## Supplemental for "In Vitro Evidence to Support Amphotericin B and Flucytosine Combination Therapy for Talaromycosis"

*Authors contributed equally

**Corresponding authors

**Corresponding Authors:**

**Supplementary File 1: *Talaromyces marneffei* isolates**

The clinical isolates were revived on yeast peptone dextrose (YPD) agar plates and grown in the yeast form at 37°C. After 3 to 6 days, a single colony was sub-cultured on a new YPD plate to achieve purity. Pure yeast cultures were maintained at 4°C for up to 2 weeks and sub-cultured at 37°C for 3 days for antifungal susceptibility experiments.

**Supplementary File 2: Methods for antifungal drug preparation**

Drug stocks, drug dilutions, and drug plates were prepared according to the Clinical and Laboratory Standards Institute (CLSI) guidelines [1]. AmB and 5FC (Sigma-Aldrich, St. Louis, MO, United States) were obtained as pure powders. In brief, AmB was dissolved in dimethyl sulfoxide (DMSO), and 5FC was dissolved in Roswell Park Memorial Institute (RPMI) medium (Sigma-Aldrich, United States) buffered to pH 7 with 3-(N-morpholino) propane sulfonic acid (MOPS, Sigma-Aldrich, United States). Serial dilutions of AmB and 5FC were prepared in RPMI-MOPS, and aliquots were stored at -20°C and thawed to room temperature before use. Our previous study reported MIC ranges of 0.5 – 1 µg/mL for AmB (95% inhibition) and 0.06 – 0.5 µg/mL for 5FC (50% inhibition) [2]. Based on these findings, the final drug concentration ranges tested were 0.03 – 2 μg/mL for AmB and 0.004 – 2 µg/mL for 5FC.

**Supplementary File 3: Antifungal susceptibility testing by our validated CLSI-based colorimetric assay**

In brief, on the day of inoculation, *T. marneffei* yeast cells were harvested from the YPD plates and suspended in phosphate-buffered saline (PBS). Using an Ultrospec^®^ cell density meter (Biochrom, Holliston, MA, United States), the inoculum was standardized to an optical density at 600 nm (OD_600nm_) range of 0.52 – 0.58, which is equivalent to 1 – 5 x 10^6^ CFU/mL. The inoculum was further diluted in RPMI-MOPS, which resulted in a working inoculum of 1 – 5 x 10^3^ CFU/mL. A total of 100µL of inoculum was added to each well of the prepared drug plate, excluding the negative control well. The plates were sealed and incubated at 37°C. After 24 hours of incubation, alamarBlue was added to all wells, and the plates were incubated at 37°C for an additional 48 hours. Following incubation, FI was measured at an excitation of 570 nm and emission of 590 nm using a fluorescence spectrophotometer (BMG LABTECH, Cary, NC, United States). The percentage reduction of fungal growth was calculated for each well in reference to the positive and negative control well.

**Supplementary File 4:** **Equation A for calculating the fractional inhibitory concentration index (FICI) by checkerboard assay**

$$\left( \mathbf{A} \right) FICI= \frac{{MIC}_{amphotericin B in combination}}{{MIC}_{amphotericin B alone}}+ \frac{{MIC}_{flucytosine in combination}}{{MIC}_{flucytosine alone}}$$

**Supplementary File 5: Methods for time-kill assay**

In brief, a standardized *T. marneffei* yeast inoculum of 10^5^ cells/mL was prepared with PBS. The MIC ranges of AmB and 5FC were determined using our CLSI-based colorimetric broth microdilution method [2]. Dose-response curves were generated using AmB at 0.25, 0.5, 1, and 2 times the MIC, and for 5FC at 0.25, 0.5, 1, 5, 10, and 20 times the MIC, reflecting clinically achievable drug levels in plasma [3,4]. For AmB and 5FC combination testing, sub-MIC concentrations of AmB (0.25 and 0.5 times the MIC) were used, as these were concentrations that were not rapidly fungicidal (as determined by the time-kill experiment for AmB alone) and thus permitted accurate assessment of the added effect of 5FC, which was tested at concentrations 1, 5, 10, and 20 times the MIC. Ten-milliliter tubes were prepared with the fungal suspension, antifungal drugs (alone or in combination), and RPMI-MOPS. A control tube containing no drug was included in all experiments. The tubes were incubated at 37°C at 250 rpm. At each pre-determined timepoint (24, 48, 72, 96, and 120 hours), 0.1 – 0.3 mL was aliquoted from each testing condition, diluted appropriately, and plated on YPD in duplicate. Fungal growth was measured by the colony forming units (CFUs/mL).

**Supplementary File 6: Quality control**

Drug concentrations were standardized based on the expected MIC of the *Candida krusei* ATCC 6258 reference strain, according to CLSI methods. *C. krusei* ATCC 6258 was included as a quality control in all susceptibility experiments, alongside *T. marneffei* strain 11CN-20-091 as an internal control. On each day of inoculation, the final *T. marneffei* inoculum was grown on YPD plates at 37°C for 5 days to check the viability of the inoculum used. A CFU count within the range of 5 – 30 was considered optimal.

**Supplementary File 7:** **Table 1 shows the distribution of the minimum inhibitory concentrations and combination effect between amphotericin B and flucytosine for 60 *Talaromyces marneffei* clinical isolates.**

| **Interaction Type** | **MIC_95_ (µg/mL)** | | **FIC _AmB_** | **MIC_95_ (µg/mL)** | | | **FIC _5FC_** | **FICI**  FIC_AmB_+FIC_5FC_ |
| --- | --- | --- | --- | --- | --- | --- | --- | --- |
|  | **AmB** | **AmB+5FC** |  | **5FC** | **5FC+AmB** | |  |  |
| ***Synergy*** *(FICI ≤ 0.5), n = 4* | | | | | | | | |
| 11CN-21-012 | 2.00 | 0.50 | 0.25 | 1.00 | | 0.06 | 0.06 | 0.31 |
| 11CN-21-040 | 1.00 | 0.25 | 0.25 | 0.13 | | 0.02 | 0.13 | 0.38 |
| 11CN-27-012 | 2.00 | 0.50 | 0.25 | 0.50 | | 0.06 | 0.13 | 0.38 |
| 11CN-27-017 | 2.00 | 0.50 | 0.25 | 0.25 | | 0.06 | 0.25 | 0.50 |
| ***Indifference*** *(0.5 < FICI ≤ 4), n = 56* | | | | | | | | |
| 11CN-03-002 | 0.50 | 0.25 | 0.50 | 0.50 | | 0.13 | 0.25 | 0.75 |
| 11CN-03-006 | 1.00 | 0.50 | 0.50 | 0.25 | | 0.00 | 0.02 | 0.52 |
| 11CN-03-007 | 0.25 | 0.02 | 0.06 | 0.25 | | 0.25 | 1.00 | 1.06 |
| 11CN-03-008 | 0.25 | 0.03 | 0.13 | 0.13 | | 0.13 | 1.00 | 1.13 |
| 11CN-03-009 | 0.50 | 0.50 | 1.00 | 0.25 | | 0.02 | 0.06 | 1.06 |
| 11CN-03-014 | 0.25 | 0.13 | 0.50 | 0.50 | | 0.13 | 0.25 | 0.75 |
| 11CN-03-015 | 0.50 | 0.13 | 0.25 | 1.00 | | 0.50 | 0.50 | 0.75 |
| 11CN-03-021 | 0.50 | 0.02 | 0.03 | 0.50 | | 0.25 | 0.50 | 0.53 |
| 11CN-03-029 | 1.00 | 0.50 | 0.50 | 0.50 | | 0.06 | 0.13 | 0.63 |
| 11CN-03-037 | 0.25 | 0.13 | 0.50 | 0.25 | | 0.13 | 0.50 | 1.00 |
| 11CN-03-039 | 0.25 | 0.13 | 0.50 | 1.00 | | 0.50 | 0.50 | 1.00 |
| 11CN-03-040 | 0.25 | 0.06 | 0.25 | 1.00 | | 0.50 | 0.50 | 0.75 |
| 11CN-03-048 | 0.25 | 0.13 | 0.50 | 1.00 | | 0.50 | 0.50 | 1.00 |
| 11CN-03-051 | 0.50 | 0.25 | 0.50 | 0.25 | | 0.13 | 0.50 | 1.00 |
| 11CN-03-068 | 1.00 | 1.00 | 1.00 | 0.25 | | 0.00 | 0.02 | 1.02 |
| 11CN-03-070 | 1.00 | 0.50 | 0.50 | 0.25 | | 0.06 | 0.25 | 0.75 |
| 11CN-03-078 | 0.25 | 0.25 | 1.00 | 0.50 | | 0.02 | 0.03 | 1.03 |
| 11CN-03-083 | 0.50 | 0.25 | 0.50 | 0.25 | | 0.06 | 0.25 | 0.75 |
| 11CN-03-086 | 1.00 | 0.50 | 0.50 | 0.25 | | 0.03 | 0.13 | 0.63 |
| 11CN-03-098 | 2.00 | 1.00 | 0.50 | 0.50 | | 0.01 | 0.02 | 0.52 |
| 11CN-03-104 | 0.50 | 0.25 | 0.50 | 0.50 | | 0.25 | 0.50 | 1.00 |
| 11CN-03-108 | 1.00 | 1.00 | 1.00 | 0.13 | | 0.00 | 0.03 | 1.03 |
| 11CN-03-116 | 1.00 | 0.50 | 0.50 | 0.25 | | 0.13 | 0.50 | 1.00 |
| 11CN-03-120 | 1.00 | 0.50 | 0.50 | 0.25 | | 0.06 | 0.25 | 0.75 |
| 11CN-03-121 | 1.00 | 0.06 | 0.06 | 0.25 | | 0.13 | 0.50 | 0.56 |
| 11CN-03-129 | 1.00 | 0.50 | 0.50 | 0.25 | | 0.06 | 0.25 | 0.75 |
| 11CN-03-130 | 0.50 | 0.25 | 0.50 | 0.50 | | 0.25 | 0.50 | 1.00 |
| 11CN-03-140 | 1.00 | 0.50 | 0.50 | 0.13 | | 0.06 | 0.50 | 1.00 |
| 11CN-03-147 | 1.00 | 0.06 | 0.06 | 0.50 | | 0.25 | 0.50 | 0.56 |
| 11CN-03-148 | 1.00 | 0.25 | 0.25 | 0.06 | | 0.03 | 0.50 | 0.75 |
| 11CN-03-153 | 1.00 | 0.50 | 0.50 | 0.25 | | 0.00 | 0.02 | 0.52 |
| 11CN-03-154 | 0.50 | 0.25 | 0.50 | 0.06 | | 0.03 | 0.50 | 1.00 |
| 11CN-03-158 | 1.00 | 0.50 | 0.50 | 0.50 | | 0.03 | 0.06 | 0.56 |
| 11CN-20-002 | 1.00 | 0.50 | 0.50 | 0.25 | | 0.06 | 0.25 | 0.75 |
| 11CN-20-005 | 0.50 | 0.25 | 0.50 | 0.50 | | 0.25 | 0.50 | 1.00 |
| 11CN-20-008 | 1.00 | 0.50 | 0.50 | 0.50 | | 0.25 | 0.50 | 1.00 |
| 11CN-20-016 | 1.00 | 0.03 | 0.03 | 0.25 | | 0.13 | 0.50 | 0.53 |
| 11CN-20-023 | 0.50 | 0.25 | 0.50 | 0.13 | | 0.06 | 0.50 | 1.00 |
| 11CN-20-029 | 0.50 | 0.02 | 0.03 | 0.13 | | 0.13 | 1.00 | 1.03 |
| 11CN-20-044 | 0.50 | 0.25 | 0.50 | 0.13 | | 0.03 | 0.25 | 0.75 |
| 11CN-20-052 | 1.00 | 0.50 | 0.50 | 2.00 | | 0.06 | 0.03 | 0.53 |
| 11CN-20-065 | 0.50 | 0.50 | 1.00 | 0.25 | | 0.02 | 0.06 | 1.06 |
| 11CN-20-068 | 1.00 | 0.50 | 0.50 | 0.25 | | 0.06 | 0.25 | 0.75 |
| 11CN-20-070 | 1.00 | 0.50 | 0.50 | 1.00 | | 0.13 | 0.13 | 0.63 |
| 11CN-20-078 | 1.00 | 0.03 | 0.03 | 0.50 | | 0.25 | 0.50 | 0.53 |
| 11CN-20-079 | 0.50 | 0.25 | 0.50 | 0.06 | | 0.03 | 0.50 | 1.00 |
| 11CN-20-091 | 0.25 | 0.13 | 0.50 | 0.25 | | 0.03 | 0.13 | 0.63 |
| 11CN-20-102 | 1.00 | 0.50 | 0.50 | 0.25 | | 0.13 | 0.50 | 1.00 |
| 11CN-20-120 | 1.00 | 0.50 | 0.50 | 0.25 | | 0.03 | 0.13 | 0.63 |
| 11CN-26-003 | 0.50 | 0.13 | 0.25 | 0.06 | | 0.03 | 0.50 | 0.75 |
| 11CN-21-013 | 2.00 | 1.00 | 0.50 | 0.13 | | 0.00 | 0.03 | 0.53 |
| 11CN-21-022 | 1.00 | 0.50 | 0.50 | 0.25 | | 0.03 | 0.13 | 0.63 |
| 11CN-21-028 | 2.00 | 1.00 | 0.50 | 1.00 | | 0.02 | 0.02 | 0.52 |
| 11CN-21-041 | 0.25 | 0.13 | 0.50 | 0.06 | | 0.02 | 0.25 | 0.75 |
| 11CN-26-024 | 0.50 | 0.25 | 0.50 | 0.06 | | 0.02 | 0.25 | 0.75 |
| 11CN-27-009 | 0.25 | 0.13 | 0.50 | 0.06 | | 0.03 | 0.50 | 1.00 |
| Mean | 0.68 | 0.24 | 0.45 | 0.28 | | 0.06 | 0.33 | 0.77 |
|  | (95% CI: 0.58-0.80) | (95% CI: 0.19-0.32) | (0.23) | (95% CI: 0.22-0.34) | | (95% CI: 0.04-0.08) | (0.25) | (0.22) |
| Mode | 1 | 0.5 | 0.5 | 0.25 | | 0.06 | 0.5 | 0.75 |

The MICs of AmB and 5FC were defined as the lowest drug concentration that resulted in at least 95% inhibition of fungal growth. The geometric means with the 95% confidence intervals are reported for the MICs of AmB and 5FC, alone, and in combination. The arithmetic means with the standard deviations are reported for FICs of AmB and 5FC, and the FICI for 60 isolates.

Abbreviations: 5FC, flucytosine; 95% CI, 95% confidence interval; AmB, amphotericin B; FIC, fractional inhibitory concentration; FICI, fractional inhibitory concentration index; MIC, minimum inhibitory concentration.
